# PTSD blood transcriptome mega-analysis: Shared inflammatory pathways across biological sex and modes of traumas

**DOI:** 10.1101/123182

**Authors:** Michael S. Breen, Daniel S. Tylee, Adam X. Maihofer, Thomas C. Neylan, Divya Mehta, Elisabeth Binder, Sharon D. Chandler, Jonathan L. Hess, William S. Kremen, Victoria B. Risbrough, Christopher H. Woelk, Dewleen G. Baker, Caroline M. Nievergelt, Ming T. Tsuang, Joseph D. Buxbaum, Stephen J. Glatt

## Abstract

Transcriptome-wide screens of peripheral blood during the onset and development of posttraumatic stress disorder (PTSD) indicate widespread immune dysregulation. However, little is known as to whether biological sex and the type of traumatic event influence shared or distinct biological pathways in PTSD. We performed a combined analysis of five independent PTSD blood transcriptome studies covering seven types of trauma in 229 PTSD and 311 comparison individuals to synthesize the extant data. Analyses by trauma type revealed a clear pattern of PTSD gene expression signatures distinguishing interpersonal (IP)-related traumas from combat-related traumas. Co-expression network analyses integrated all data and identified distinct gene expression perturbations across sex and modes of trauma in PTSD, including one wound-healing module down-regulated in men exposed to combat traumas, one IL12-mediated signaling module up-regulated in men exposed to IP-related traumas, and two modules associated with lipid metabolism and MAPK-activity up-regulated in women exposed to IP-related traumas. Remarkably, a high degree of sharing of transcriptional dysregulation across sex and modes of trauma in PTSD was also observed converging on common signaling cascades, including cytokine, innate immune and type I interferon pathways. Collectively, these findings provide a broad view of immune dysregulation in PTSD and demonstrate inflammatory pathways of molecular convergence and specificity, which may inform mechanisms and diagnostic biomarkers for the disorder.

## INTRODUCTION

Posttraumatic stress disorder (PTSD) is a debilitating disorder that develops after exposure to a traumatic event and increases vulnerability to adverse health outcomes. The estimated lifetime prevalence of PTSD is ~5-6% in men and ~10-12% in women (Kessler, 1995), and even higher among recent war-veterans with estimates as high as ~20% (Ramchand *et al*, 2010). Although extensive work has identified putative risk factors that are associated with PTSD (DiGanji *et al*, 2013), the identification of discrete diagnostic biomarkers for the disorder remains elusive. Heterogeneity in susceptibility to PTSD suggests that the response of an individual to trauma may depend on biological sex as well as the type of adverse event (*e.g.* early life adversity, violence, assault, accidents, combat). These factors, in turn, may determine downstream consequences, such as perturbation of biological pathways, making it unlikely that a valid, singular biomarker will be specific to all PTSD cases.

Research into the mechanisms underlying the onset and development of PTSD converge on hypothalamic-pituitary-adrenal (HPA) axis and immune system functioning (Daskalakis *et al*, 2016; Cohen *et al*, 2016). As such, several studies have examined pro-inflammatory cytokines and glucocorticoid activity in peripheral blood mononuclear cells (PBMC) or lymphocytes in PTSD cases to build more effective models for identifying molecular factors underlying PTSD. These studies were reviewed by Passos *et al* (2016), who summarized that increases in C-reactive protein (*CRP*), interleukin 6 (*IL-6*), tumor necrosis factor alpha (*TNF-α*), interleukin 1 beta (*IL-1β*) and interferon gamma (*IFN-γ*) all underlie the onset and emergence of PTSD symptoms. However, despite the evident effects of biological sex on incidence rates of PTSD, very few of the reviewed studies examined sex differences in psychobiological or inflammatory responses to trauma. Moreover, the majority of these reports centered their analysis around pre-determined targets, limiting the ability to identify novel genes and molecular pathways relevant to the pathophysiology of PTSD.

Transcriptome-wide screens of peripheral immune cells from individuals with PTSD have extended findings from candidate gene studies through systems-wide exploration of immune system dysregulation in response to PTSD. Segman *et al* (2005) first reported on transcriptomic differences in peripheral blood from trauma survivors with PTSD on the day of emergency room visit and four months later, implicating dysregulation of transcriptional enhancers and immune activating genes. We later described blood-based transcriptomic signatures implicating sex differences in cytokine pathways activated in PTSD from a population of individuals exposed to various traumatic backgrounds (Neylan *et al*, 2011). In a separate and non-overlapping study, we also identified transcriptomic differences in central nervous system development and immune tolerance induction pathways in PTSD cases, both with and without a history of childhood maltreatment, implicating trauma-related dimorphism (Mehta *et al*, 2013). Parallel lines of research have identified candidate prognostic and diagnostic blood-based gene expression classifiers in war-veterans with PTSD (Glatt *et al*, 2015; Tylee *et al* 2015), that were largely associated with dysregulated innate immune function both prior to and following PTSD development (Breen *et al*, 2015). Collectively, while these data-driven approaches have formed the foundation of ongoing work to build PTSD biomarkers, they have yet to be widely replicated. Inconsistences may be attributable to various clinical factors (*e.g.* comorbidity, medication) and technical factors (*e.g.* various technologies and statistical methods used to evaluate data). More importantly, these studies are often severely underpowered and model gene expression changes in the context of an explicitly defined biological sex or trauma type. As such, a critical remaining question is how biological pathways in peripheral blood overlap across sex and modes of trauma in PTSD, and how this information may inform the search for more verifiable diagnostic biomarkers for the disorder.

The primary goal of the current investigation was to synthesize the existing data from transcriptome-wide gene expression studies in PTSD and to clarify their relevance to PTSD pathophysiology, while explicitly modeling sex- and trauma-related differences. To do so, we performed a mega-analysis of five independent transcriptome-wide peripheral blood studies covering seven types of trauma in 229 PTSD and 311 comparison individuals. To address our goals, a standardized multistep analytic approach was used that we have reviewed in the context of blood-based biomarker discovery in neuropsychiatric disorders (Breen *et al*, 2016), and that we have also applied to other transcriptome-wide mega-analyses (Hess *et al*, 2016; Tylee *et al*, 2016). To this end, our analyses specifically sought to: (1) determine the relatedness of PTSD gene expression signatures across different types of trauma; (2) identify candidate genes, pathways and co-regulatory networks in PTSD and determine if such alterations are distinct between different biological sexes and trauma types; and (3) construct diagnostic blood-based gene expression classifiers to differentiate PTSD cases from trauma-exposed control individuals and clarify the potential clinical utility of peripheral blood gene expression.

## METHODS

### Literature search and study criteria

To systematically identify relevant studies for our combined mega-analysis, we performed a literature search (SCOPUS) and microarray database searches (NCBI GEO and EMBL-EBI ArrayExpress) for transcriptome-wide studies of whole blood- or leukocyte-based gene expression in PTSD. Studies were included if they met the following criteria: (i) crosssectional post-trauma studies published between 2005-2015; (ii) contained individuals meeting structured diagnostic criteria for PTSD (*e.g.* DSM, PCL); (iii) also contained a trauma-exposed healthy control group. Studies were excluded on the bases of: (i) using qualitative real-time PCR or immunoassays as a means to investigate a targeted panel of candidate genes; (ii) investigating mechanisms in lymphoblastoid cell lines, skin fibroblast cultures, serum, and plasma; and (iii) secondary data integration analyses and review papers were also excluded. Five studies met these criteria for which raw gene expression data and clinical covariates (age, sex, ethnicity and trauma type) were available, and one additional study from which data were unavailable (Sarapas *et al*, 2011). Using these five studies, seven trauma-specific case-control bio-sets were curated by parsing individuals provided by Neylan *et al*, 2011 into two separate sub-groups (that is, combat- and assault-related traumas) and Mehta *et al*, 2013 into two separate sub-groups (that is, childhood- and interpersonal-related traumas). All data were obtained from either the corresponding authors of the original studies or from publicly available data repositories. For diagnostic criteria, we relied on those used at each study site, some of which were based on clinician assessments and others based on standardized screening tool. There was no additional filtering of subjects based on medical comorbidities beyond what was described on the original studies.

### Gene expression data processing and quality control

All statistical analyses were conducted in the statistical package *R.* Data from each study were processed, normalized and quality treated independently (see **Supplementary Figure 1** for workflow). Briefly, when multiple microarray probes mapped to the same HGNC symbol, the probe with the highest average expression across all samples was used for further analysis. Normalized data were inspected for outlying samples using unsupervised hierarchical clustering of samples and principal component analysis to identify potential outliers outside two standard deviations from these grand averages. Combat batch correction (Leek *et al*, 2015) was applied to remove systematic sources of variability other than case/control status, such as technical, clinical, or demographic factors both within each study (as necessary), and then across all studies using common gene symbols, forming the bases for subsequent mega-analytic case-control groups. The frequencies of circulating immune cells were estimated for each individual in each study using Cibersort cell type de-convolution (https://cibersort.stanford.edu/) (Newman *et al*, 2015). See **Supplementary File** for full details on these data processing steps.

### Differential gene expression analyses

*Trauma-specific analyses:* Differential gene expression (DGE) was performed independently for each of the seven case-control groups using the limma package (Ritchie *et al*, 2015) to detect relationships between diagnostic status and gene expression levels. The covariates age and sex were included in all models to adjust for their potential confounding influence on gene expression between main group effects. To determine the relatedness of DGE signatures across the seven trauma-specific groups, each gene list was converted into a matrix of binary gene presence/absence calls with respect to each group and a Jaccard coefficient was applied to create a gene-based phylogeny, as previously described (Diaz-Beltran *et al*, 2016).

*Mega-analytic analyses:* To increase statistical power and test for biological sex- and trauma-specific gene signatures, individual samples from different experiments were combined based on *i)* biological sex and *ii)* type of trauma to form three large mega-analytic case-control groups. First, common gene symbols across all available samples were identified (*n*_*genes*_=4,062). Then, Combat batch correction (Leek *et al*, 2015) was applied to remove systematic sources of variability other than case/control status, such as technical factors (*e.g.* difference technologies, duration of time from PTSD onset to blood acquisition), clinical factors (*e.g.* comorbidities) or demographic factors (*e.g.* ethnicity) across all individual studies. Finally, DGE analysis was performed for each mega-analytic case-control group controlling for effects of age, and unless otherwise specified, the significance threshold was a nominal *P*-value <0.05. This nominally significant *P*-value was used to yield a reasonable number of genes to include within functional annotation and gene network analyses.

### Weighted gene co-expression network analysis

Weighted gene co-expression network analysis (WGCNA) (Langfelder *et al*, 2008) was used to build signed co-expression networks using a total of 4,062 genes found in common across all experiments (see **Supplementary Information** for details). Two broad analyses were performed. First, a series of module preservation analyses sought to determine whether PTSD development influences the underlying gene co-regulatory patterns, as being preserved or disrupted, compared to controls, and *vice versa.* For these analyses, module preservation was assessed using a permutation-based preservation statistic, *Z_summary_*, implemented within WGCNA with 500 random permutations of the data (Langfelder *et al*, 2011). *Z_summary_* takes into account the overlap in module membership as well as the density and connectivity patterns of genes within modules. A *Z_summary_* score <2 indicates no evidence of preservation, *2<Z_summary_<10* implies weak preservation and *Z_summary_* >10 suggests strong preservation. Second, to increase confidence and power to detect biologically meaningful modules, a consensus network was built use all available samples. Once modules were identified from the consensus network, modules were assessed for significant associations to PTSD diagnostic status, sex and mode of trauma. Singular value decomposition of each module’s expression matrix was performed and the resulting module eigengene (ME), equivalent to the first principal component, was used to represent the overall expression profiles for each module. Differential analyses of MEs was performed using Bayes ANOVA (Kayala *et al*, 2012) (parameters: conf=12, bayes=1, winSize=5), comparing between diagnostic status, sex and mode of trauma, correcting *P*-values for multiple comparisons with *post hoc* Tukey tests.

### Functional annotation and protein interaction networks

The ToppFunn module of ToppGene Suite software (Chen *et al*, 2015) was used to assess enrichment of Gene Ontology (GO) terms using a one-tailed hyper geometric distribution with family-wise false discovery rate (FDR) at 5%. GO semantic similarity analysis was used to assess shared/unique gene content amongst GO terms using the GoSemSim semantic similarity R package (Yu *et al*, 2015), and default semantic contribution factors (‘is_a’ relationship: 0.8 and ‘part_of’ relationship: 0.5). Second, gene modules were tested for over-representation of PTSD genome-wide association study (GWAS) signatures obtained from the DisGenNet database (Pinero *et al*, 2015), retrieved using the disease-term query ‘PTSD’. Third, DGE signatures were used to build direct protein-protein interaction (PPI) networks, which can reveal key genes/transcription factors mediating the regulation of multiple target genes. PPIs were obtained from the STRING database (Franceschini *et al*, 2012) with a signature query of DGE lists from the mega-analytic case-control comparisons. We used a combined STRING score of >0.4 (*i.e.* medium-to-high confidence interactions). For visualization, the STRING network was imported into CytoScape (Shannon *et al*, 2003).

### Construction of PTSD blood-based diagnostic classifiers

BRB-Array Tools supervised classification methods (Simon *et al*, 2007) were used to construct gene expression classifiers. Three models were specified to distinguish PTSD cases from controls relative to: (1) men exposed to combat trauma (2) men exposed to IP traumas, and (3) women exposed to IP traumas. Each model consisted of three steps. First, to ensure a fair comparison, all genes in the training data with *P* <0.05 were subjected to classifier construction, respective for each mega-analytic case-control group. This heuristic rule of thumb approach was used to cast a wide net to catch all potentially informative genes, while false-positives would be pared off by subsequent optimization and cross-validation steps. Second, classifiers composed of different numbers of genes were constructed by recursive feature elimination (RFE). RFE provided feature selection, model fitting and performance evaluation via identifying the optimal number of features with maximum predictive accuracy. Third, the ability for RFE to predict group outcome was assessed by support vector machines (SVM) and compared to four different multivariate classification methods (*i.e.* diagonal linear discriminant analysis (DLDA), nearest centroid (NC), first-nearest neighbors (1NN), three-nearest neighbors (3NN)). For each of the three models, classification accuracies are reported for both the training data (70% of data) and the completely withheld test data (30% of data) as area under the receiver operating curve (AUC).

### Statistical power and sample size computation

We estimated the expected discovery rate (EDR), a multi-test equivalent to power, and sample size at a fixed number of biological replicates (n) and type I error rate (α) using the PowerAtlas software (Page *et al*, 2006). This sample size calculation method is based on studies of the distribution of *P*-values from DGE analyses from microarray studies controlling for EDR. For evaluating which *n* is best suited for a future study, we set the average probability of detecting an effect (EDR) to be >0.8 and α=0.05.

### Code and data availability

Computational code and quality controlled gene expression data are available upon request to the corresponding author and can also be directly downloaded at https://github.com/BreenMS/PTSD-blood-transcriptome-mega-analysis.

## RESULTS

### Literature search and data curation

A total of five cross-sectional PTSD studies met our criteria (Methods) for which raw gene expression data and clinical covariates were available (**Table 1**). From these five studies, seven trauma-specific case-control groups were derived, including three groups exposed to combat traumas, one group exposed to assault traumas, one group exposed to childhood-related traumas, one group exposed to emergency room (ER) accident-related traumas, and one group exposed to ‘other’ interpersonal (IP)-related traumas which could not be explicitly defined. These seven trauma-specific groups were later combined to form three large mega-analytic case-control groups, aimed at explicitly modeling for sex- and trauma-related differences (**Table 1,** bottom), including: *i)* men exposed to combat traumas (*n*_PTSD_=85, *n*_Control_=84, *k*_genes_ =10,112); *ii)* men exposed to IP traumas (*n*_Ptsd_=45, *n*_Control_=67, *k*_genes_=4,378); and *iii)* women exposed to IP traumas (*n*_PTSD_=99, *n*_Control_ =160, *k*_genes_ =4,378).

**Table 1.** Blood-based transcriptome-wide studies of Posttraumatic Stress Disorder included in the mega-analysis.

| Study | Data Type | PTSD (n) | Controls (n) | % Female | Age (Years) | Sample Type | Predominant Ancestry, % | Genes Analyzed | Additional Sample Information<sup>†</sup> |
| --- | --- | --- | --- | --- | --- | --- | --- | --- | --- |
| Breen *et al.*, 2015<br>GSE64814 | Poly-A enriched, 50bp paired-end sequencing on Illumina Hi-Seq 2000. | 46 Combat | 46 Combat | 0% | 23.3 ± 3.22 | Isolated Leukocytes | European, 55.5% | 13,944 | Marine Resilience Study (MRS) sample of U.S. Marines who served a 7 month combat deployment. Blood was drawn 3-months post-deployment for each participant. PTSD symptoms were assessed at this time using CAPS and diagnosis was determined using DSM-IV criteria for partial or full PTSD. |
| Tylee *et al.*, 2015<br>GSE63878 | Affymetrix Hu-Gene 1.0 ST Array | 24 Combat | 24 Combat | 0% | 22.2 ± 3.1 | Isolated Leukocytes | European, 72.0% | 22,772 | MRS sample of U.S. Marines who served 7-9 month combat deployment. Blood was drawn 3-months post-deployment for each participant. PTSD symptoms were assessed and diagnostic status was determined using the CAPS. |
| Mehta *et al.*, 2013 | Illumina HT-12 version 3.0 | 54 Childhood<br>67 Other IP | 38 Childhood<br>166 Other IP | 73.6% | 42.0 ± 12.6 | Whole Blood | African American, 88.9% | 9,193 | General medical community-based sample (Atlanta, Georgia) selected for traumatic exposure. Blood samples were obtained years after trauma exposure. PTSD symptoms were assessed using the PSS and diagnosis was determined by applying DSM-IV criteria to PSS items; control subjects were negative for current or lifetime PTSD. |
| Neylan *et al.*, 2011 | CodeLink Human Whole Genome BioArrays | 15 Combat<br>14 Assault | 14 Combat<br>15 Assault | 26.9% | 30.0 ± 6.0 | Isolated CD14+ Monocytes | European, 56.7% | 17,988 | Recruited through Veterans Affairs Medical Center PTSD Outpatient Program (San Francisco, California) and through community fliers. Included both assault and combat-related traumas. Blood samples were obtained years after trauma exposure. Symptoms were assessed and diagnostic status was determined using the CAPS. |
| Segman *et al.*, 2005 | Affymetrix Human Genome-U95A | 9 ER-trauma | 8 ER-trauma | 37.5% | 31.1 ± 11.4 | Isolated Mononuclear Cells | Jewish Ancestry, 100% | 9,668 | Acute trauma exposure in emergency room setting. Samples collected at 4 months after trauma onset. PTSD cases met DSM-IV criteria for PTSD 4 months post-trauma. |
| Total: 5 studies |  | 229 | 311 |  |  |  |  |  |  |
| Combat-Trauma, Men | Mega-Analytical Group 1 | 85 | 84 | 0% | 24.4 ± 4.7 | Combined | European, 63.6 % | 10,112 |  |
| IP-Trauma, Men | Mega-Analytical Group 2 | 45 | 67 | 0% | 41.1 ± 12.8 | Combined | African American 69.7% | 4,378 |  |
| IP-Trauma, Women | Mega-Analytical Group 3 | 99 | 160 | 100% | 39.5 ± 12.3 | Combined | African American 76.1% | 4,378 |  |

We attempted to include available information pertaining to each sample, with particular emphasis on the type of trauma exposure, the time point of biological sample acquisition, and the diagnostic criteria used for each study. All details were obtained from the referenced articles. Abbreviations: Clinician administered PTSD Scale (CAPS), cluster of differentiation 14-positive (CD14+), Diagnostic and Statistical Manual of Mental Disorders-IV (DSM-IV), quality control-passing (QC+), interpersonal trauma (IP).

### Between-trauma comparisons

Following standardized data pre-processing procedures (see Methods and Supplementary File), the proportions of circulating immune cells were estimated for all individuals since complete cell counts with leukocyte differentials were not available. Comparative analyses of the estimated immune cell type proportions showed no significant differences between PTSD cases and controls in any of the seven trauma-specific case-control groups (**Supplementary Table 1**), suggesting that cell type frequencies would not confound downstream analyses. Subsequently, to determine the overall relatedness of the trauma-specific groups, seven lists of covariate adjusted differential gene expression (DGE) signatures were generated, then converted into a binary matrix of gene presence/absence calls. Distance-based clustering with pairwise similarity was measured via Jaccard coefficient. A high degree of DGE similarity formed two distinct branches that clustered the five IP trauma groups how from the three combat trauma groups (**Figure 1A**). Pair-wise overlaps of the seven DGE lists were used to further quantify this result (**Figure 1B-C**) and identified a number of significant overlaps between: childhood and assault traumas (⋂=35, *P*=0.05); childhood and ER-related traumas (⋂=31, *P*=0.03); assault and ER-related traumas (⋂=18, *P*=0.04); ER-related traumas and IP-related traumas (⋂=9, *P*=0.03); and combat traumas between Breen *et al.*, and Neylan *et al.*, (⋂=23, *P*=6.9e-9) (**Supplementary Table 2**). No genes were consistently differentially expressed across all five IP trauma groups (**Figure 1B**), although two genes showed consistent but weak effects across all three combat trauma groups (**Figure 1C**); interferon induced protein 44 like (*IFI44L*) was over-expressed while G protein subunit gamma 11 (*GNG11*) was underexpressed in PTSD cases relative to controls. Notably, no gene in any of the above comparisons survived FDR *P* <0.05.

### Mega-analytic comparisons

To increase statistical power, these seven trauma-specific case-control groups were combined to form three large mega-analytic case-control groups designed to explicitly model for differences in sex and modes of trauma (that is, combat and IP traumas) and DGE lists were generated for each comparison (**Supplementary Table 3**). Comparatively equal numbers of over- and under-expressed genes were observed in men exposed to combat traumas (*n*_*up*_=150, *n_down_=*174) and men exposed to IP traumas (*n*_*up*_=114, *n*_*down*_=145), while women exposed to IP traumas displayed significantly more overexpressed genes than under-expressed genes (*n*_*up*_ =123, *n_down_*=63, *P*=1.2e-05; two-tailed proportions test) (**Figure 2A**). DGE indicated small, but significant, gene overlaps between men exposed to combat traumas and women exposed to IP traumas (⋂=18, *P*=1.1e-08), men exposed to combat traumas and men exposed to IP traumas (⋂=7, *P*=0.04), and between men and women exposed to IP traumas (⋂=15, *P*=2.3e-08) (**Figure 2B**). Notably, one gene, interferon induced protein with tetratricopeptide repeats 3 (*IFIT3*), was significantly differentially expressed in all comparisons, albeit with different directions of change. Overall, DGE signatures were associated with PTSD diagnosis and not with any other factors, including age, ancestry, study site or estimated cell-type frequencies (**Supplementary Figure 2**).

Though few genes were differentially expressed in all comparisons, functional annotation of these DGE signatures indicated a high degree of overlap of commonly perturbed biological processes between men exposed to combat- and IP-related traumas (*n*=27, *P*=2.1e-14), men exposed to combat traumas and women exposed to IP traumas (*n*=16, *P*=3.3e-12), and between men and women exposed to IP-related traumas (*n*=42, *P*=1.3e-137) (**Figure 2C**). In addition to several common biological processes, numerous unique gene-sets were also identified for each comparison (**Supplementary Table 4**) suggesting that differences in sex and trauma types may impact distinct biological processes. However, in exploring the semantic similarity between these distinct gene-sets, we identified a series of relevant, biologically meaningful interactions, positioning each distinct biological process as a component of a broad ‘stress response system’ (**Figure 2D**). To support this observation, we tested whether candidate genes that are dysregulated together indeed interact with each other at the protein level. A significant overrepresentation of direct protein interactions was identified for each DGE list, and a union of all three networks was constructed (**Supplementary Figure 3**). Notably, *IFIT3* demonstrated protein-level interactions with partners across all three networks, among other interferon proteins such as *IFI44L, IRF7, IFI44, IFI35, IFIT4* and *IRF4.* The network generated from men exposed to combat traumas featured several genes with a high degree of connectivity involved in type I interferon signaling and antiviral responses, including *DDX58, IFIH1, IFIT1/2, MX1, RSAD2, STAT1*, and members of the *OAS* gene family. Comparably, genes related to men with a history of IP trauma formed a relatively unique network with the most highly connected genes included *EZR, H2AFZ, IMPDH2, JUND, STAT5B*, and *SYK.* The network generated for women with a history of IP trauma, consisted of *ABL1, ATM, TNF* and *UBB*, which demonstrated the highest degree of connectivity.

Markedly, of the 15 biological processes found at the intersection of all comparisons (**Figure 2C**), all terms were strongly enriched for innate immune responses, cytokine signaling, and cytokine production (**Figure 3A**). Surprisingly, the genes within each of these gene-sets were predominantly over-expressed among men exposed to combat traumas and women exposed to IP traumas, but were under-expressed in men exposed to IP traumas. To further quantify this observation, the concordance of transcriptome-wide DGE patterns was calculated among the three mega-analytic case-control comparisons first constraining to all genes specific to innate immune or cytokine signaling and then using all remaining genes (**Figure 3B-D**). Positive associations in changes of innate immune and cytokine genes were observed between men exposed to combat traumas and women exposed to IP traumas (*r*=0.61, *P*=3.3e-34), and negative associations were observed between men exposed to combat and IP traumas (*r*=−0.37, *P*=6.4e-12), and between men and women exposed to IP traumas (*r*=−0.29, *P*=1.2e-12). Next, we sought to determine whether these biological processes were specific to PTSD or found in common with other neuropsychiatric disorders including major depression, schizophrenia, bipolar disorder and autism spectrum disorder, by implementing series of cross-disorder overlap comparisons, at both the individual gene and gene-ontology level (**Supplementary Figure 4, Supplementary Table 4**). Indeed, the majority of innate immune and cytokine signatures were more strongly related to a universal diagnosis of PTSD across differences in biological sex and modes of traumas, rather than in these other disorders.

**Figure 4.**
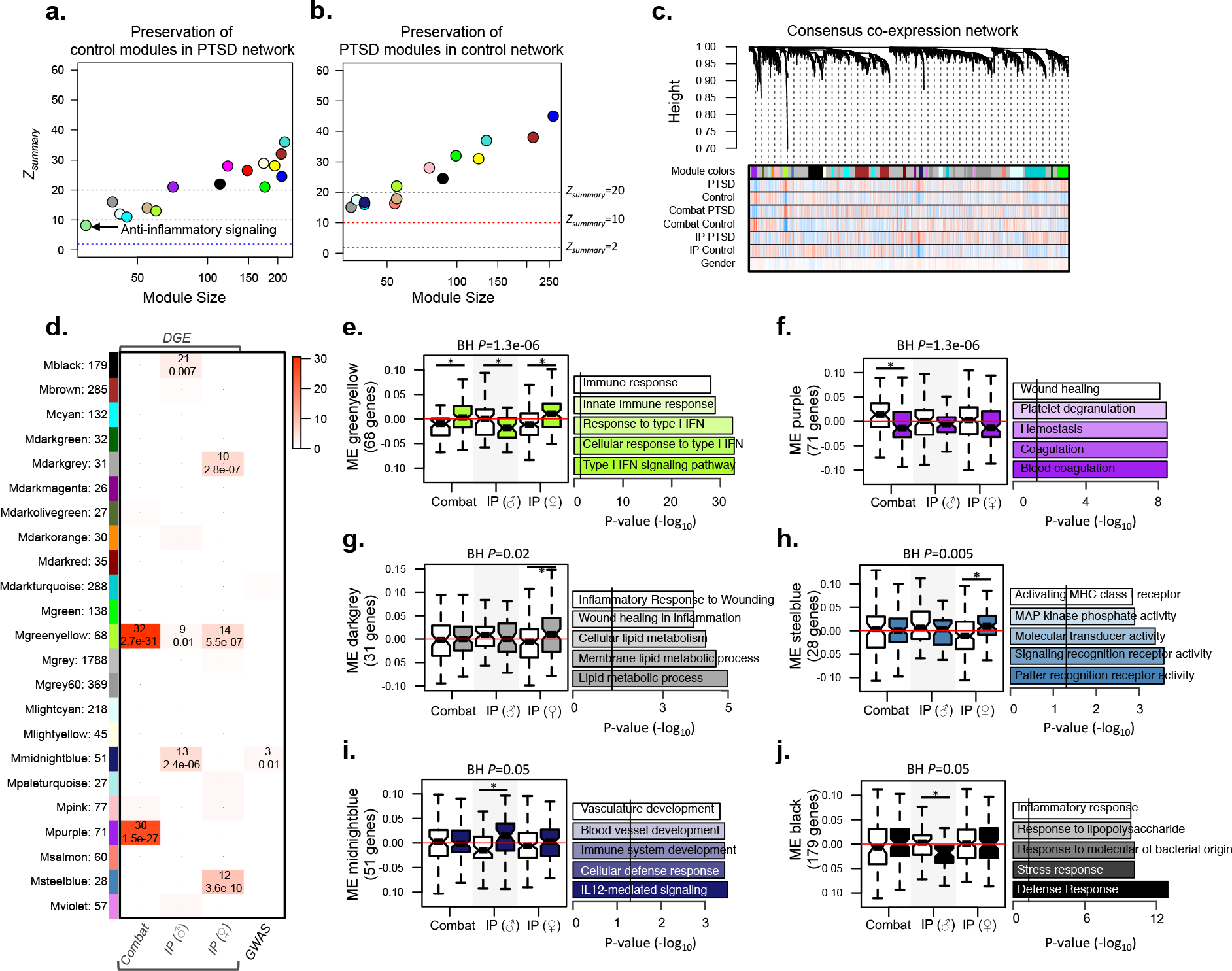
Consensus weighted gene co-expression network analysis (WGCNA). **(A)** Module preservation identified one module in control individuals with weak preservation in PTSD cases. **(B)** All PTSD modules were preserved in control samples. (C) Hierarchical clustering tree (dendrogram) of the consensus network and all samples comprising 4,062 genes. Each line represents a gene (leaf) and each low-hanging cluster represents a group of co-expressed genes with similar network connections (branch) on the tree. The first band underneath the tree indicates the twenty-three detected modules and subsequent bands indicate gene-trait correlation when red indicates a strong relationship and blue indicates a strong negative relationship. **(D)** Gene modules were enriched for DGE signatures and PTSD GWAS signatures curated from DisGenNet database (x-axis). The number of genes within each colored gene module are depicted (y-axis). The top number in each cell indicates the number of genes overlapping and the bottom number indicates the *P*-value significance of overlap using the Fisher’s Exact Test. **(E-J)** A Bayes ANOVA was used on ME values to test for significance between case-control status across different biological sex and trauma types and (*) indicates BH P<0.05, implying significant PTSD differences. For each module the top five most significant biological processes and/or pathways are reported.

### Stratified gene co-expression module preservation analyses

WGCNA was used to assess the extent of module preservation by integrating all PTSD cases compared to all control individuals using a permutation-based preservation statistic (*Z_summary_*, see Methods). This analysis identifies large differences in gene co-regulatory patterns as being disrupted in PTSD cases relative to controls, or *vise versa* (**Figure 4A-B**). In control individuals, sixteen modules were identified and one module implicated in antiinflammatory signaling was weakly preserved (*Z_summary_*=8.6) in PTSD cases, including genes *IL10, TNFSF14* and *LILRB1.* In the reverse approach, fourteen modules were identified across PTSD cases and all were highly preserved within control individuals, indicating that major changes in the underlying gene-gene connectivity may not be a basis for the pathology of PTSD. A separate series of between trauma-type comparisons were also performed (see **Supplementary File** and **Supplementary Figure 5** for details).

### Consensus gene co-expression network analyses

Subsequently, we used the higher confidence and completeness of a consensus network by combining all individuals across the three mega-analytic case-control groups (**Figure 4C**). This analysis identified 23 co-expression modules, which were tested for enrichment of DGE signatures and PTSD-related GWAS signals (**Figure 4D**). Module eigenvalues (MEs) were then subjected to a Bayes ANOVA testing to compare the extent of module expression differences between diagnostic status, sex and type of trauma (**Figure 4E-J**). A greenyellow module (68 genes) implicated in type I interferon-mediated signaling cascades and enriched with differentially expressed genes from all three mega-analytic comparisons was significantly over-expressed in PTSD-affected men exposed to combat traumas and PTSD-affected women exposed to IP traumas, but was under-expressed in PTSD-affected men exposed to IP traumas. A purple module (71 genes) implicated in blood coagulation and wound healing was under-expressed in PTSD-affected men exposed to combat traumas. A darkgrey module (31 genes) enriched with inflammatory response to wounding and cellular lipid membrane metabolic process, and a steelblue module (28 genes) implicated in intracellular pattern recognition receptor signaling and mitogen-activated protein kinase (MAPK) phosphate activity were both over expressed among PTSD-affected women exposed to IP traumas. A midnight blue module (51 genes) enriched for IL12-mediated signaling and vascular development was over-expressed among PTSD-affected men exposed to IP traumas. This module also contained three potential PTSD risk-related genes (*FASLG, IFN-γ, RORA*) previously identified through GWAS, reflecting greater than chance overlap (*P*=0.012). A black module (179 genes) implicated in immune response and response to bacterial lipopolysaccharide was under-expressed among PTSD-affected men exposed to IP traumas.

### PTSD blood-based diagnostic classifier evaluation

Supervised class prediction methods were used to directly assess the putative clinical utility of blood-based gene expression measurements for objective PTSD diagnosis, as well as to identify any discriminant gene(s) that may have been overlooked in our previous analyses. Three models were specified to distinguish PTSD cases from control individuals for each mega-analytic group. Classification accuracies were reported on the training data (70% of data) as well as independently withheld test data (30% of data), and support vector machine (SVM) learning consistently outperformed the other methods **(Supplementary Figure 6)**. First, when distinguishing between PTSD cases and control individuals exposed to combat traumas, classification accuracies reached 81% in the training data and 60% in the withheld test data when the expression of 40 genes was used with SVM classification. Second, when separating PTSD-affected men from controls individuals exposed to IP traumas, classification accuracy reached 77% in the training data and 65% on the withheld test data when the expression of 60 genes was used with SVM. Third, when separating PTSD-affected women from controls individuals exposed to IP traumas, classification accuracy reached 67% in the training data and 58% on the withheld test data when the expression of 25 genes as used with SVM; two common genes were selected by both IP-related trauma models across men and women (*GNB5* and *DGCR14*). Further details regarding these analyses are provided in Methods and **Supplementary Table 6.**

### Statistical power and sample size estimation

To inform the design of future cross-sectional studies in PTSD, we estimated the expected discovery rate (EDR; a multi-test equivalent to power) and sample size for each mega-analytic case-control group using lists of *P*-values derived from DGE analyses. Effect sizes estimated from these data were assumed to be fixed with a nominal error rate α=0.05 and several different sample sizes (n) were evaluated. To determine how many case and control samples need to be included in a future study, we set the threshold of inclusion to EDR >0.8. Overall, the sample size needed to reach a power of 0.8 for men exposed to IP trauma was *n* of 700, for women exposed to IP trauma was *n* of 7000, while men exposed to combat trauma was *n* of 10,000 (**Supplementary Table 7**).

## DISCUSSION

To our knowledge, this is the largest transcriptome-wide analysis of PTSD conducted to date. Our combined mega-analysis covered 229 PTSD cases and 311 comparison individuals, enabling us to increase statistical power and to explicitly test whether differences in sex and trauma type play a role in perturbing common or distinct molecular pathways in PTSD. A battery of statistical tests were applied and we report several findings. First, re-analyses of seven trauma-specific case-control groups revealed a high degree of relatedness amongst IP-related traumas separate from combat-related traumas. Second, once individual samples were combined to form three mega-analytic case-control groups, we observed unique PTSD DGE signatures in men exposed to combat traumas, men exposed to IP traumas and women exposed to IP traumas, which all converged on shared biological processes of innate immune, cytokine and type I interferon signaling. Third, stratified network analysis identified low module preservation between control individuals exposed to different traumas, but high module preservation between PTSD cases exposed to the same types of trauma, suggesting that the underlying molecular response to different trauma types may be more homogenous in PTSD cases. Fourth, one anti-inflammatory module within control individuals was weakly preserved across all PTSD cases, indicating the disruption/absence of anti-inflammatory gene co-regulation within PTSD cases. Fifth, upon integrating all data to construct one consensus network, numerous sex and trauma-specific modules were identified, and one module implicated in innate immunity and type I interferon signaling was significantly associated to PTSD in all three mega-analytic groups. Sixth, supervised multivariate classification methods constructed diagnostic PTSD classifiers for each mega-analytic group with moderate-to-low classification accuracies on withheld test data. Taken together, our analyses indicate that while small-to-moderate effect sizes are the standard for cross-sectional post-trauma studies of PTSD, our findings consistently converge on similar down-stream inflammatory pathways irrespective of sex and the type of traumatic event.

A novel finding was the small, but significant, between-trauma overlap of DGE signatures indicating the existence of a trauma-specific and across-trauma convergent gene regulation and signaling (**Figure 1**). Indeed, we previously tested the hypothesis that differences in trauma may impact the stress response in PTSD and we identified distinct gene expression signatures between PTSD cases with and without a history of childhood maltreatment that also converged on similar cellular processes (Mehta *et al*, 2013). Here, we extend upon these results and position them among a broad background of individuals exposed to a range of different traumatic events. Notably, not one of the seven trauma-specific case-control comparisons resulted in a gene passing FDR *P*-value <0.05, indicating the underpowered nature of the observed data. Thus, to increase statistical power, this initial result provided enough empirical support to combine the data to form three mega-analytic case-control groups, enabling us to explicitly model for differences in sex and trauma.

**Figure 1.**
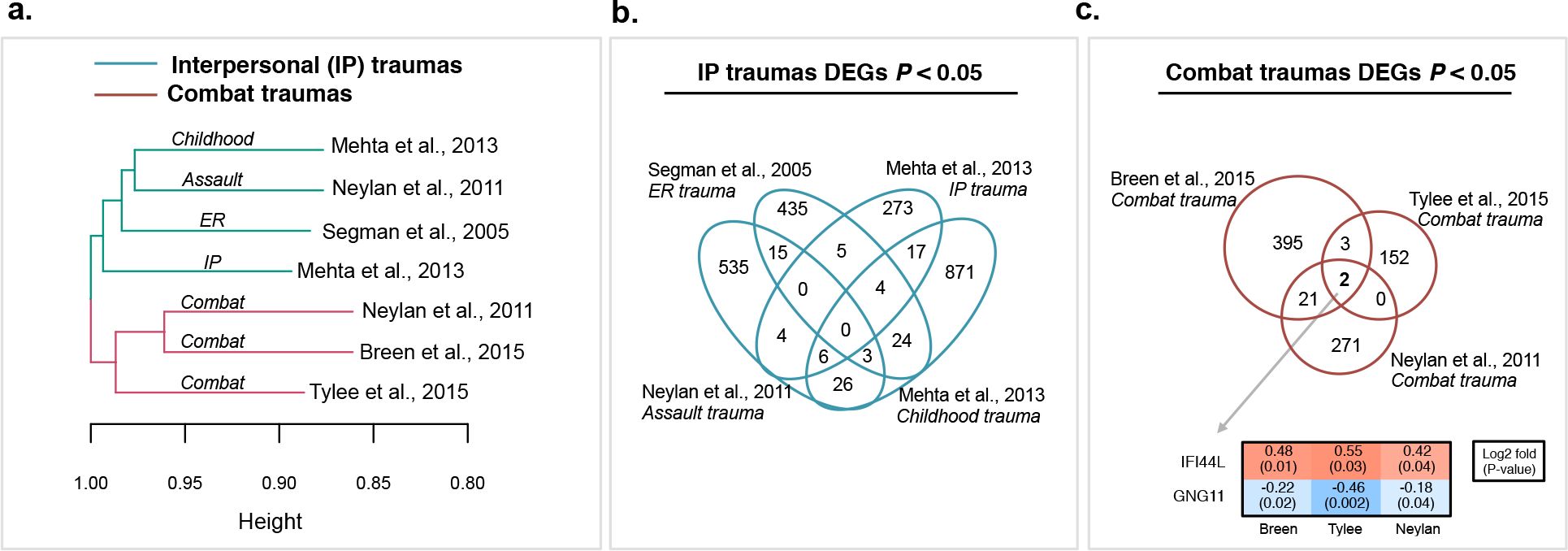
Concordance of differential gene expression (DGE) analyses across seven trauma-specific case-control groups. **(A)** Jaccard clustering of PTSD DGE signatures from the seven trauma-specific case-control groups. Overlap of PTSD DGE signatures found in common across **(B)** interpersonal (IP) traumas and (C) combat traumas. For combat-related traumas, interferon Induced Protein 44 Like (*IFI44L)* was consistently over-expressed and G Protein Subunit Gamma 11 (*GNG11*) was consistently under-expressed in PTSD cases relative to control individuals. All analyses were adjusted for age and sex. Supplementary Table 2 contains full lists of overlapping gene symbols.

Our central finding was the identification of largely unique DGE perturbations specific to each mega-analytic case-control group, which converged on common biological processes of innate immune, cytokine and type I interferon signaling cascades (**Figure 2**). Our consensus network approach validated and refined this result through identification of a discrete co-regulated gene module (68 genes) implicated in innate immune and type I interferon signaling that displayed divergent expression patterns across sex and traumas in PTSD (**Figure 4E-J**). This result was previously reported in PTSD cases exposed to combat trauma (Breen *et al*, 2015), and is now replicated across a larger sample. With respect to potential mechanisms, some of the earliest observed effects of inflammatory cytokines in PTSD underline their impact on the HPA-axis, via negative feedback regulation (Michopoulos *et al*, 2015). Enhanced negative feedback regulation of the HPA axis function is a hallmark of PTSD and is reflected by increased responsiveness to glucocorticoids as manifested by decreased cortisol concentrations following dexamethasone administration (Michopoulos *et al*, 2015; Yehuda *et al*, 2002). Inflammatory cytokines have also been shown to access the brain and interact with virtually every pathophysiological domain relevant to PTSD, including neurotransmitter metabolism, neuroendocrine function and neural plasticity (Felger *et al*, 2013). In doing so, peripheral cytokine signals activate relevant brain cell types that serve to amplify central inflammatory responses and conserved behavioral responses. Notably, we also found evidence for biological processes significantly overrepresented in only one of the three mega-analytic groups, suggesting that sex and type of trauma may influence different molecular pathways in PTSD (**Figure 2C**). However, in determining the relatedness between each of these distinct biological responses, we found that all dysregulated biological processes interacted and collectively formed a biologically meaningful ‘stress response system’ (**Figure 2D**). Indeed, a dysfunctional HPA axis-immune interface has been previously associated with similar immune and metabolic disturbances, including cell cycle, altered cytokine balance, blood coagulation and lipid and metabolic processes (Silverman *et al*, 2013; Silverman *et al*, 2012; Elenkov *et al*, 2000). Under this standard, each dysregulated biological process represents one piece of a larger stress response system, irrespective of sex and trauma, that ultimately converges on shared inflammatory pathways.

**Figure 2.**
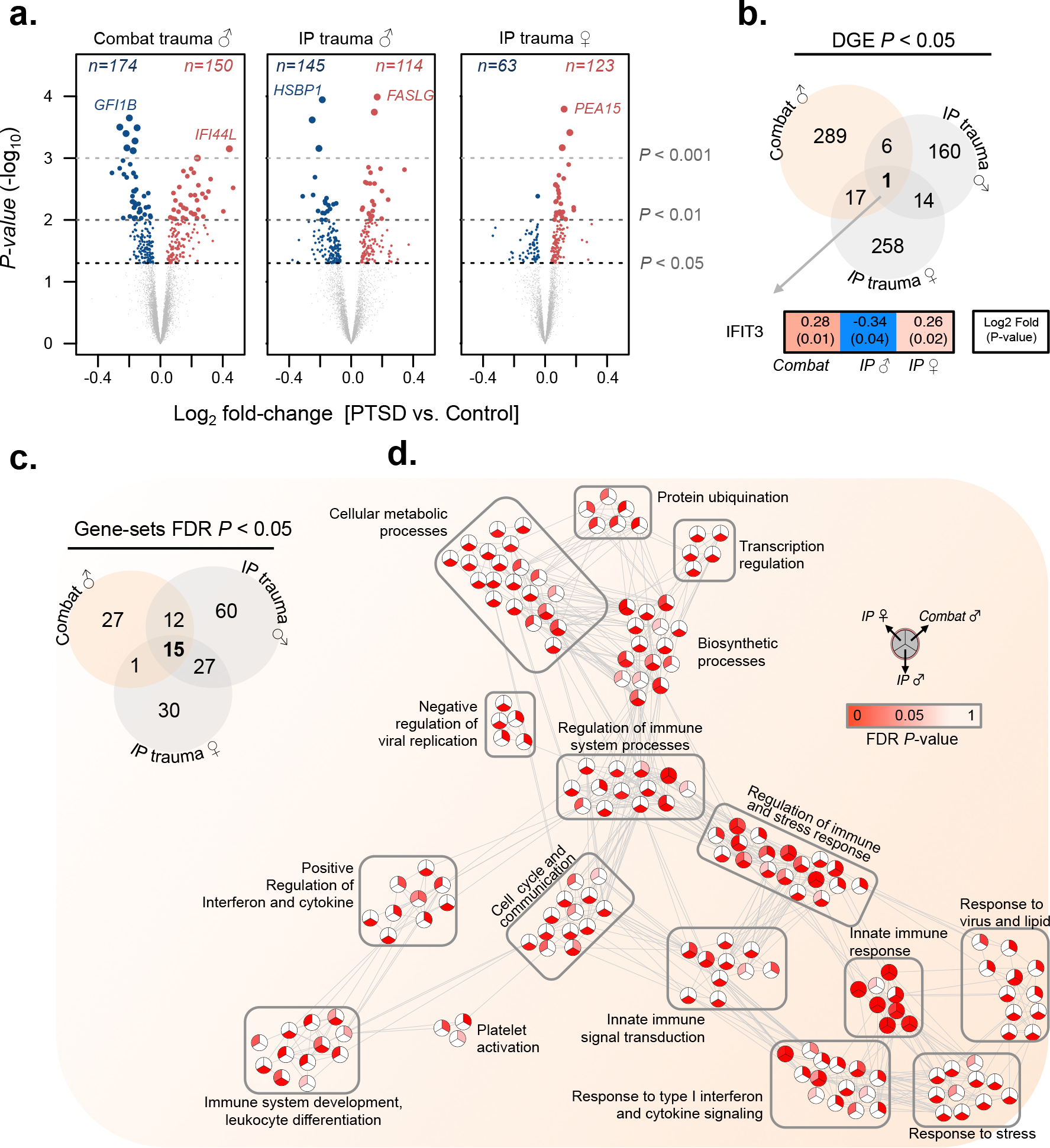
Differential gene expression (DGE) and gene ontology (GO) analyses for the three mega-analytic case-control comparisons. **(A)** Volcano plots compare the extent of log_2_ fold-change and -logi_0_ *P*-value significance for DGE signatures in all three comparisons. The top over and under expressed genes are labeled for each comparison. Overlap of **(B)** significant DGE signatures and (C) enriched GO gene-sets for all comparisons are displayed. **(D)** Relatedness of all significantly dysregulated gene-sets by semantic similarity. Nodes represent GO-terms and edges represent semantic similarity > 0.5 (high degree of gene overlap). Nodes are split into thirds and shaded by FDR *P*-value significance for each mega-analytic group.

Some of the observed PTSD-related effects involved lower expression levels for cytokine-related genes, specifically in men exposed to IP trauma compared to women, and men exposed to combat trauma (**Figure 3**). Differences in inflammatory cytokine levels in trauma survivors with PTSD have previously been reported (Gill *et al*, 2009). Discrepancies may potentially reflect different degrees of HPA-axis response suppression to glucocorticoid activation (Freidenberg *et al*, 2010), or alternatively, differences associated with the duration since the traumatic experience as well as other confounding factors. One study examining the levels of inflammatory markers within a refugee population with PTSD postulated that differences could partly be explained by a variable environmental component associated with less antigen exposure (Söndergaard *et al*, 2004). We also acknowledge that gene expression perturbations may be fundamentally different in controls with prolonged exposure to conflict zones compared to controls exposed to IP traumas, which may be influencing these results. To address this issue, a series of pair-wise coexpression preservation analyses were performed using matched controls exposed to different types of trauma (**Supplementary Figure 5**). These analyses indicated that gene co-expression modules involved in processes other than inflammatory signatures differed on the basis of exposure to different traumatic events.

**Figure 3.**
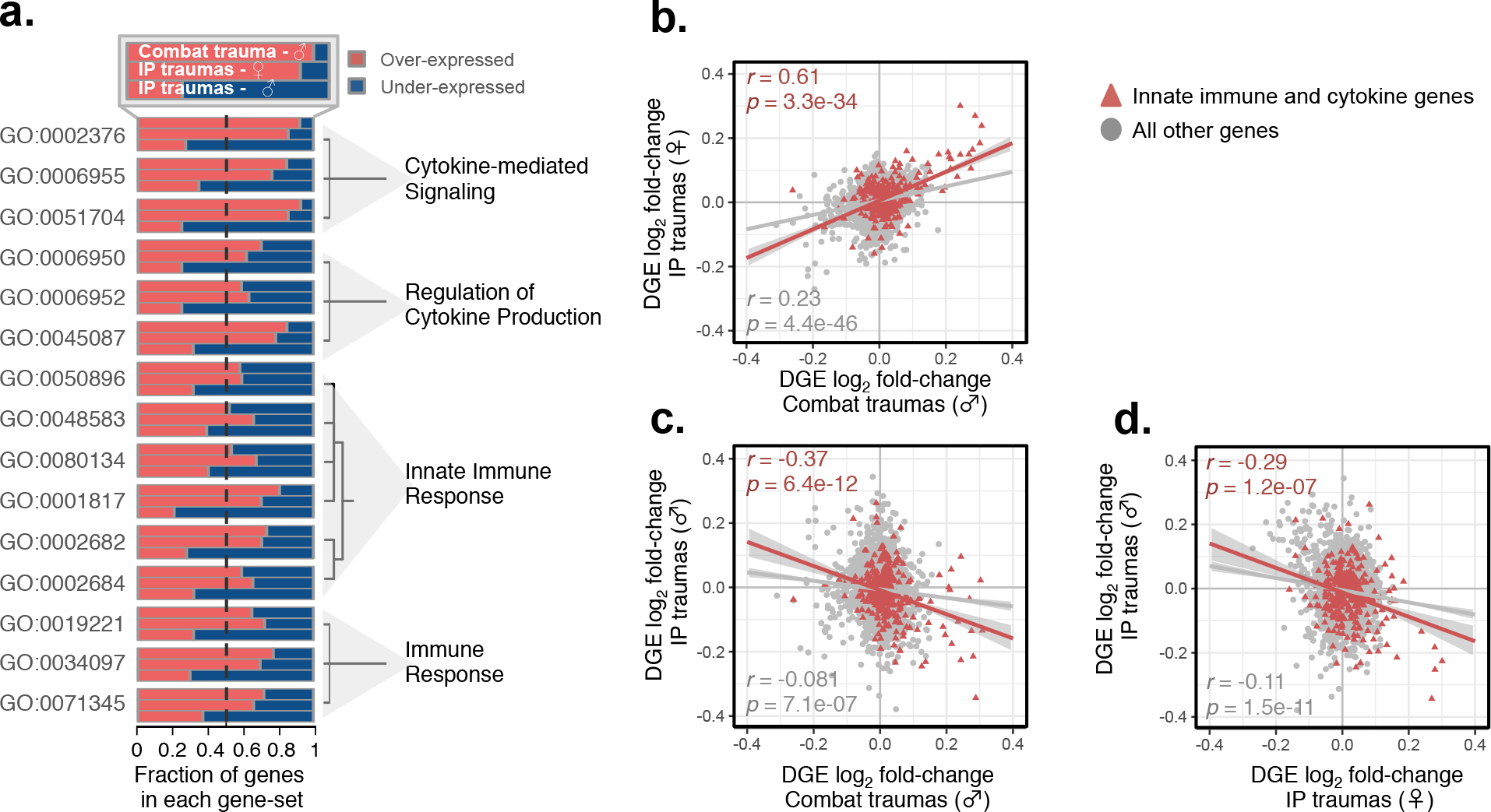
Concordance of transcriptome-wide innate immune and cytokine signatures. **(A)** Semantic similarity for all fifteen common GO term pairs (left) clustered by hierarchical clustering method with the fraction of over- and under-expressed genes for each GO term (middle) as well as common ancestors (right). Log_2_ fold-changes (PTSD vs. controls) were used in a series of pair-wise correlations amongst the three mega-analytic case-control groups and focused on innate immune and cytokine genes relative to all other genes. **(B)** A positive association between PTSD-affected men exposed to combat traumas and PTSD-affected women exposed to IP traumas. Negative associations between PTSD-affected men exposed to IP traumas and (C) PTSD-affected men exposed to combat traumas and **(D)** PTSD-affected women exposed to IP traumas. Pearson correlation coefficients were applied.

Distinct gene expression perturbations were also identified, including *(i)* one wound healing module down-regulated in men with a history of combat trauma, *(ii)* two modules implicated in lipid processes and MAPK-activity up-regulated in women exposed to IP-related trauma and *(iii)* one IL12-mediated signaling module up-regulated in men exposed to IP-related trauma. Regarding to the first distinction *(i)*, research highlights the role of platelets in innate and adaptive immune responses and suggest that platelet activation and reactivity is dysregulated by mental stress. A stress response involving blood platelets has also been shown to be a critical biomarker of hemostatic, thrombotic, and inflammatory perturbations (Pacák *et al*, 2001). Notably, this result is also a re-affirmation of our previous finding indicating decreased wound healing and blood coagulation in war-veterans with PTSD (Breen *et al*, 2015), now replicated in a larger cohort of samples. Regarding the second distinction *(ii)*, the MAPK pathway functions as a mediator of cellular stress, including inflammation, by also modulating the levels of glucocorticoid receptor phosphorylation, ultimately leading to differences in cellular transcriptional activity (Galliher-Beckley *et al*, 2011; Reul, 2014). Since signaling cascades, such as the MAPK, couple to numerous receptors for stress-related neurotransmitters and neuropeptides (Whitaker *et al*, 2014), future work should determine more precisely which neurotransmitter signaling systems are driving the observed traumatic stress- and sex-induced changes. Regarding the final distinction *(iii)*, the IL-12 signaling pathway initiates innate and adaptive immune responses in part by promoting NK cell toxicity as well as the differentiation of naive CD4+ T cells into T helper 1 cells, and induces the production of IFN-γ, which is also a member of this gene module. Here, up-regulation of IL-12 signaling in men exposed to IP trauma indicates immune priming that may promote inflammation. With respect to sex dimorphism, increased percentage of IL-12 producing monocytes and lymphocytes in response to physiological concentrations of testosterone has been reported in men compared to women (Posma *et al*, 2014). Taken together, these distinct biological perturbations all align with a common pro-inflammatory pathology across sex and modes of trauma in PTSD. However, a great deal of research is needed to further delineate the precise mechanisms, as well as cause and effect relationships, underlying these inflammatory signatures and sex disparities.

Our study also has several limitations. First, sample size and power estimates indicated that our enhanced sample size is still underpowered and a substantially larger number of biological replicates are needed (**Supplementary Table 7**). These estimates are echoed by our inability to identify multiple test corrected DGE signatures and to construct accurate diagnostic blood-based PTSD classifiers, all of which reported low-to-moderate classification accuracies on withheld test data. Second, it is likely that our results may be influenced by clinical heterogeneity (e.g. medical comorbidity, medication) among PTSD cases, potentially contributing to the diminished power. While our effort to carefully measure the contribution of potential confounding factors on our gene expression results demonstrated that DGE signatures were associated with PTSD diagnosis and not with any other factors (*e.g.* age, ancestry, study site, cell-type frequencies), other unmeasured factors may also influence the results. Additionally, to fully understand the contribution of ancestry to gene expression variation, future studies may consider integrating principal components of ancestry from paired GWAS data if available. Third, we were unable to distinguish biological sex differences for combat-related traumas due to a large ascertainment bias of men with a history of combat exposure. Fourth, though our estimates of cell type fractions implied no differences between case-control groups, we are unable to determine gene expression changes specific to any particular cell type. Finally, as the combined data are cross-sectional with considerable variation in the amount of time from PTSD onset to blood sample acquisition, we are unable to determine whether expression differences represent expression signature of past or current PTSD, or it is a marker of pretrauma vulnerability to PTSD development. Similarly, some traumas are defined by event (*e.g.* combat, assault) but others are defined by time (*e.g.* childhood) or place (*e.g.* ER). Likewise, both childhood and ER traumas might be due to similar trauma types, including assaults, which are unknown in the current study and may effect the interpretation of our results.

In sum, these data provide evidence for shared inflammatory profiles in peripheral blood gene expression across sex and modes of trauma in PTSD as evident by transcriptional dysregulation and co-expression on processes of innate immune, cytokine and type I interferon signaling. Moreover, the fact that several unique biological processes were also affected across sex and trauma types that ultimately formed components of a broader stress response system, underscore a shared underlying molecular pathology. While existing animal and cellular work support the sex-dependent effects specific to MAPK and IL-12 signaling modules, further research is needed to delineate cause and effect relationships. Collectively, these findings may have implications for identifying objective diagnostic biomarkers, disease mechanisms and therapeutic interventions in immune disturbances for PTSD.

## Acknowledgements

We gratefully acknowledge the individual volunteers and their families, as well as the Marine and Navy volunteers for their military service and for their participation in these studies. Additionally, we thank Segman and colleagues (2005), as well as others, who have placed their data in the public domain. The Marine Resiliency Study (MRS) was supported by VA Health Service Research and Development project no. SDR 09-0128, the Marine Corps, and the Navy Bureau of Medicine and Surgery (DGB) and MRS II (DGB, VBR) and its Demonstration Project (CMN) by the Naval Medical Research Center's Advanced Medical Development program (Naval Medical Logistics Command Contract #N62645-11-C-4037. MRS-II acknowledges support from the administrative core, A Patel, A De La Rosa, members of the MRS-II Team and the Veterans Medical Research Foundation (VMRF). We thank the Marine and Navy volunteers for their military service and for their participation in this study. We also thank Dr. Anna Tocheva for further critical reading/assessment of our manuscript.

## Author contributions

DST and SJG assembled all the data from independent sources. MSB designed the research question, analyzed the data and wrote the manuscript, with DST. All remaining authors contributed to the generation of the original data and provided critical reading/assessment of the current manuscript.

